# Transcriptome Analysis of Cortical Tissue Reveals Shared Sets of Down-Regulated Genes in Autism and Schizophrenia

**DOI:** 10.1101/029132

**Authors:** Shannon E. Ellis, Rebecca Panitch, Andrew B. West, Dan E. Arking

## Abstract

Autism (AUT), Schizophrenia (SCZ), and bipolar disorder (BPD) are three highly heritable neuropsychiatric conditions. Clinical similarities and genetic overlap between the three disorders have been reported; however, the causes and the downstream effects of this overlap remain elusive. By analyzing transcriptomic RNA-Sequencing data generated from post-mortem cortical brain tissues from AUT, SCZ, BPD and control subjects, we have begun to characterize the extent of gene expression overlap between these disorders. We report that the AUT and SCZ transcriptomes are significantly correlated (p<0.001), while the other two cross disorder comparisons (AUT-BPD, SCZ-BPD) are not. Among AUT and SCZ, we find that the genes differentially expressed across disorders are involved in neurotransmission and synapse regulation. Despite lack of global transcriptomic overlap across all three disorders, we highlight two genes, *IQSEC3* and *COPS7A*, which are significantly down-regulated compared to controls across all three disorders, suggesting either shared etiology or compensatory changes across these neuropsychiatric conditions. Finally, we tested for enrichment of genes differentially expressed across disorders in genetic association signals in AUT, SCZ or BPD, reporting lack of signal in any of the previously published GWAS. Together, these studies highlight the importance of examining gene expression from the primary tissue involved in neuropsychiatric conditions, cortical brain. We identify a shared role for altered neurotransmission and synapse regulation in AUT and SCZ, in addition to two genes that may more generally contribute to neurodevelopmental and neuropsychiatric conditions.

## INTRODUCTION

The aggregation of psychiatric conditions and symptoms in families has long been recognized(1-5) with more recent genetic analyses suggesting overlap between a number of disorders (1,6-9). Recent studies considering SNP-based genetic correlation demonstrated marked correlation between schizophrenia (SCZ) and bipolar disorder (BPD) and to a lesser extent between SCZ and autism spectrum disorder (ASD) (1), suggesting shared genetic etiologies. However, due to limited brain tissue availability, there have been fewer studies at the level of gene expression. We and others hypothesize that gene expression studies may begin to unravel how genetic correlations may functionally overlap in neuropsychiatric disorders.

In a recent publication, Zhao et. al suggested that SCZ and BPD show concordant differential gene expression (R=0.28) and that the genes contributing to this overlap are enriched for genetic association signal in both SCZ and BPD while highlighting several biological pathways (10). Two separate recent studies of gene expression in autism (AUT) have resolved gene expression changes related to altered synaptic and neuronal signaling as well as immunological differences in autism-affected brains (11,12). In particular, a marked increase was observed in gene expression related to alternative activation of the innate immune system, or the M2 response in autism-affected brains, relative to controls (12).

Here we set out to analyze RNA sequencing (RNA-Seq) data in combination from AUT, SCZ and BPD to identify cross-disorder transcriptomic relationships. We highlight the highly correlated nature of the SCZ and AUT transcriptomes, which together demonstrate a downregulation of genes involved in neurotransmission and synapse regulation across the two disorders.

## MATERIALS AND METHODS

### Autism Sample Information

RNA-Seq for 104 cortical brain tissue samples across three brain regions (BA10, BA19, BA44/45), comprising 57 samples from 40 control subjects and 47 samples from 32 autism (AUT) subjects was previously carried out (12). We note that, as in the initial publication of these data (12), AUT samples harboring CNVs recurrent in autism spectrum disorder have not been included in these analyses. Details related to samples, sequencing, quality control, and informatics can be found in Gupta et. al (12) and are summarized in Supplemental Table 1.

### Schizophrenia and Bipolar Disorder Sample Information

RNA-Seq data was obtained from the Stanley Medical Research Institute (SMRI, <u>http://www.stanleyresearch.org/</u>) consisting of eighty-two (31 SCZ, 25 BPD and 26 controls) anterior cingulate cortex (BA24) samples. Detailed sequencing information can be found in Zhao et. al (10). Sample information for those included in this analysis can be found in Supplemental Table 2.

### RNA-Seq, Alignment & Quality Control

Sequencing, alignment, quality control and gene expression estimation for the AUT samples were carried out as previously described (12). The reads from both the AUT and SMRI sequencing were subjected to a common analysis pipeline (12) in which quality control of raw sequences included removing both polyA stretches and adaptor sequence contamination using a Python script, ‘cutadapt’ (v1.2.1) (13). Sequences were then aligned to the Genome Reference Consortium Human build 37 (GRCh37/hg19) assembly using TopHat2 (14,15) allowing for only uniquely aligned sequences with fewer than three mismatches to align.

### Gene Expression Estimation and Normalization

Gene count estimates were obtained for 62,069 Ensembl gene annotations (GRCh37/hg19) using HTSeq (<u>http://www-huber.embl.de/users/anders/HTSeq/</u>) under an intersection-strict model. Of these, 8,856 genes with at least 10 reads across 75 percent of the SMRI samples were then normalized for gene length and GC content using Conditional Quantile Normalization (CQN) (16). In the AUT samples, the 13,262 genes previously included for analysis (12) were normalized for gene length and GC content using CQN. Outliers were then removed from the CQN normalized gene expression estimates on a per-gene basis as described previously (17). In either data set, any sample whose gene expression value was more than 2.7 standard deviations (sd) from the mean of the gene expression was excluded from analysis at that particular gene prior to linear modeling.

### Differential Gene Expression Analysis (DGEA)

Due to the unique experimental design in which multiple brain regions were sequenced from the same individual, AUT gene expression estimates were fit using a linear mixed effects model, with subject ID included as a random intercept term, and case-control status as the primary variable of interest. Age, sex, site of sample collection, brain region and twelve surrogate variables (SVs) (18) were included as fixed effects in the model to account for known and unknown covariates. SVs function to remove batch effects and sources of noise in gene expression data by adjusting for unknown or unmodeled sources of variation and are therefore included for analysis (18).

SCZ and BPD RNA-Seq data were analyzed using standard linear regression with case-control status as the primary variable of interest. The known covariates to which we had access and that were included in the analysis by Zhao et. al (10) (age, sex, cumulative antipsychotic use, brain pH, and postmortem interval (PMI)) were incorporated into the model here along with SVs to account for unknown sources of variation.

Because the SCZ and BPD cases share controls, two separate differential gene expression analyses were performed. For the comparison to AUT, all cases (SCZ or BPD) and all controls from the SMRI dataset were included in the analysis. Alternatively, when SCZ and BPD were to be compared directly, we employed a strategy similar to how these data were handled previously, in which controls were divided randomly in half (10). One set of controls was then compared to the SCZ cases while the other set of controls was compared to the BPD cases. This procedure was carried out 100 times for each cross-disorder comparison and the Z-scores (β/se) were recorded for each gene for each run. The median Z-score for each gene across these 100 runs was then used for analyses comparing SCZ to BPD.

### Null DGEA

To obtain a null set of differential gene expression values, each of the analyses in the previous section was carried out modeling the data exactly as described above save for the permutation of case-control status. In AUT datasets, case-control status was randomized between samples from the same collection sites, as described previously (12). To minimize the possibility of reporting false-positive findings, one-thousand null permutations were utilized to determine significance.

### Calculating Genes Differentially Expressed Across Disorders

To determine which genes were differentially expressed across disorders, Z-scores were multiplied across each of the three disorder comparisons (Z_SCZ_* Z_BPD_, Z_SCZ_* Z_AUT_, Z_BPD_* Z_AUT_). Genes with large cross-disorder Z-scores were considered to be differentially expressed across disorders, with significance determined by permutation. For each cross-disorder comparison, the most extreme cross-disorder Z-score for each of these 1000 null permutations was recorded. Of these values, the cross-disorder cutoff for significance (defined at p<0.05) to determine which genes were differentially expressed across disorders was determined by taking the value for which only 5% of the null values were more extreme.

To determine differentially and concordantly expressed genes (DCEGs) common to all three disorders, Z-scores were multiplied for the 2,895 genes with Z-scores in the same direction across all three disorders (Z_AUT_* Z_SCZ_* Z_BPD_). As SCZ and BPD are directly compared in the analysis, split-control generated Z-scores for SCZ and BPD were utilized to account for the shared control samples. To assess significance, the same analysis was carried out with 1000 null permutations as described above.

### Calculating the Correlation of DCEGs Across Phenotypes

Pearson’s correlation coefficient (R) was calculated for the Z-scores from each disorder comparison (SCZ-AUT, SCZ-BPD, BPD-AUT) to assess the similarity of genes differentially expressed across disorders. To determine the significance of this correlation, Pearson’s correlation coefficient was calculated after testing each of the 1000 null permutations.

### Pathway Analysis of DCEGs

Pathway enrichment analysis was carried out on genes differentially expressed across disorders. GO gene sets were downloaded from MsigDB (1466 gene sets, <u>http://www.broadinstitute.Org/gsea/msigdb/collections.jsp#C5</u>). For each gene and across all three disease comparisons, Z-scores were summed across disorders using Stouffer’s method (19) and pathways were tested for enrichment (details can be found in Supplemental Methods). Significance was determined empirically by permutation for each cross-disorder comparison (1.51×10^-4^ for AUT-SCZ, 1.72×10^-4^ for AUT-BPD, and 4.25×10^-6^ for SCZ-BPD).

As a complementary approach, we utilized two open source programs for pathway analysis: WebGestalt (v2, <u>http://bioinfo.vanderbilt.edu/webgestalt/</u>) (20,21) to run a Gene Ontology (GO) analysis (22,23) and DAVID(24) (v6.7, <u>https://david.ncifcrf.gov/</u>) for functional pathway analysis. As the input for these approaches requires gene lists, we input genes that were differentially expressed (absolute value(Z-score) > 2.2) in both disorders of the comparison: 1) SCZ-AUT (191 genes), 2) BPD-AUT (38 genes), and 3) SCZ-BPD (16 genes).

GO analysis used a hypergeometric test for enrichment utilizing the Benjamini-Hochberg method (25) for multiple test correction. GO categories whose adjusted p-values < 0.001 were considered to be statistically significantly enriched. For DAVID, gene lists were uploaded and a ‘Functional Annotation Chart’ was generated using default settings. Functional categories whose Bonferroni-adjusted p-value<0.05 were reported as significant.

To ensure that results from these analyses were not biased by the different number of genes input into the pathway analysis, we also carried out the GO and DAVID analyses described above with a fixed number of 191 genes from each cross-disorder comparison.

### Enrichment for Genetic Signal Analysis

GWAS results were downloaded from the Psychiatric Genetic Consortium (PGC, <u>http://www.med.unc.edu/pgc/</u>) for autism, bipolar disorder, and schizophrenia. Gene-based p-values were computed on the summary data for each disorder using FAST (v1.8) (26) for the 8,856 genes included in the cross disorder DGEA. Details for settings used can be found in the Supplemental Methods. To test for enrichment of genetic signal, we first took suggestive genes (gene-based p<0.05) for each individual GWAS (SCZ, BPD, and AUT) and compared these to p-values from the DGEA. Data were plotted in a QQ-plot among 100 null permutations to look for enrichment relative to the null data. To ensure that this analysis was not a reflection of the gene-based p-value restriction imposed on the data, a more permissive (p<0.1) and more restrictive (p<0.01) GWAS cutoff were used and the same enrichment analysis carried out.

### Code availability

Code used throughout for data processing, quality control, and analysis are available from corresponding author.

## RESULTS

### Sample Summary

Of the 105 samples in the SMRI array collection, 82 cortical brain samples (BA24) were sequenced and included for analysis (31 SCZ, 25 BPD, and 26 controls). To accompany these data, 104 AUT samples from three cortical brain regions (BA10, BA19, BA44/45) were included for analysis, composed of 57 control and 47 AUT samples. A summary of sample statistics are provided in Table 1 with detailed sample information in Supplemental Tables 1 and 2 for AUT and SMRI data, respectively. Further sample information can be found in the original publications (10,12).

**Table 1.** Sample Summary. Abbreviations: AUT, autism; BPD, bipolar disorder; SCZ, schizophrenia; CTL, control

|  | N | Unique<br>Individuals | Mean<br>Age<br>(years) | Sex |  |
| --- | --- | --- | --- | --- | --- |
|  |  |  |  | F | M |
| **AUT** |  |  |  |  |  |
| CTL | 57 | 40 | 20 | 12 | 33 |
| AUT | 47 | 32 | 24 | 9 | 18 |
| TOTAL | 104 | 72 | 22 | 21 | 51 |
| **SMRI** |  |  |  |  |  |
| CTL | 26 | 26 | 44 | 4 | 22 |
| BPD | 25 | 25 | 47 | 12 | 13 |
| SCZ | 31 | 31 | 42 | 7 | 24 |
| TOTAL | 82 | 82 | 44 | 23 | 59 |

### Genes Differentially Expressed Across SCZ, BPD, and AUT

Nine genes were differentially expressed (p<0.05) in both SCZ and AUT. None were significant when comparing BPD to SCZ, and one gene reached significance in the AUT-BPD comparison (Table 2, Supplemental Table 3, and Supplemental Table 4). We note that the single gene differentially expressed between AUT-BPD, *IQSEC3*, is significant in both AUT-SCZ and AUT-BPD comparisons. The relatively large Z-scores in SCZ (Z=-3.59) and BPD (Z=-3.46) suggest this result is not simply driven by the altered gene expression in AUT alone.

Differentially expressed genes (DEGs) across all three disorders were identified in a joint analysis of genes whose direction of effect was consistent across all three disorders (Z_AUT_*Z_SCZ_*Z_BPD_). Two genes, *IQSEC3* (Z=-35.45, p=0.001) and *COPS7A*(Z=-22.52, p=0.017), are transcriptome-wide significant (p<0.05, absolute value (Z_AUT_*Z_SCZ_*Z_BPD_) > 19.56), indicating a common role for altered gene expression of these genes across all three neuropsychiatric disorders (Table 2 & Supplemental Figure 1). We note that these two genes, *IQSEC3* and *COPS7A*, are syntenic (12p13.33 and 12p13.31, respectively) with their expression being markedly correlated in both the SMRI and AUT data sets (R=0.41 and R=0.70, respectively) (Supplemental Figure 2).

**Table 2.** Genes significantly differentially expressed across disorders. Abbreviations: AUT, autism; BPD, bipolar disorder; SCZ, schizophrenia; Sig., significant; chr, chromosome; Z, Z-Score

|  | # Sig. Genes | Cross-Disorder Sig. Cutoff | Ensembl Gene IDs | Gene Name | chr | Z<sub>cross-disorder</sub> | Z<sub>AUT</sub> | Z<sub>SCZ</sub> | Z<sub>BPD</sub> |
| --- | --- | --- | --- | --- | --- | --- | --- | --- | --- |
| **AUT-SCZ** | 9 | 12.42 | ENSG00000106261 | *ZKSCAN1* | 7 | 15.24 | 4.08 | 3.74 | 0.68 |
|  |  |  | ENSG00000172005 | *MAL* | 2 | 14.66 | 5.24 | 2.80 | 1.20 |
|  |  |  | ENSG00000120645 | *IQSEC3* | 12 | 14.53 | -4.04 | -3.59 | -3.46 |
|  |  |  | ENSG00000046653 | *GPM6B* | X | 14.16 | 3.62 | 3.91 | 0.26 |
|  |  |  | ENSG00000167191 | *GPRC5B* | 16 | 13.85 | 3.72 | 3.72 | 0.78 |
|  |  |  | ENSG00000129521 | *EGLN3* | 14 | 13.62 | 4.76 | 2.86 | 0.13 |
|  |  |  | ENSG00000164068 | *RNF123* | 3 | 12.81 | -3.46 | -3.71 | -0.99 |
|  |  |  | ENSG00000134780 | *DAGLA* | 11 | 12.54 | -4.22 | -2.98 | -0.57 |
|  |  |  | ENSG00000183597 | *TANGO2* | 22 | 12.53 | -3.83 | -3.27 | 0.25 |
| **AUT-BPD** | 1 | 12.29 | ENSG00000120645 | *IQSEC3* | 12 | 14.00 | -4.04 | -3.59 | -3.46 |
| **SCZ-BPD** | 0 | 21.71 | NA | -- | -- | -- | -- | -- | -- |
| **AUT-SCZ-BPD** | 2 | 19.56 | ENSG00000120645 | *IQSEC3* | 12 | -35.45 | -4.04 | -2.95 | -2.97 |
|  |  |  | ENSG00000111652 | *COPS7A* | 12 | -22.52 | -3.31 | -3.14 | -2.17 |

### Correlation in gene expression across SCZ, BPD, and AUT

The transcriptomic relationship across disorders and correlation of test-statistics (Z-scores) was investigated. SCZ-AUT demonstrated the most significant correlation (R=0.298, p<0.001). SCZ-BPD also demonstrated a positive correlation (R=0.11). This level of correlation was neither significant (p=0.41) nor as high as previously reported (R=0.28) (10). Similarly, the correlation between AUT and BPD was minimal and did not differ significantly from the null (R=0.06, p=0.25). (Figure 1, Supplemental Figure 3).

**Figure 1.**
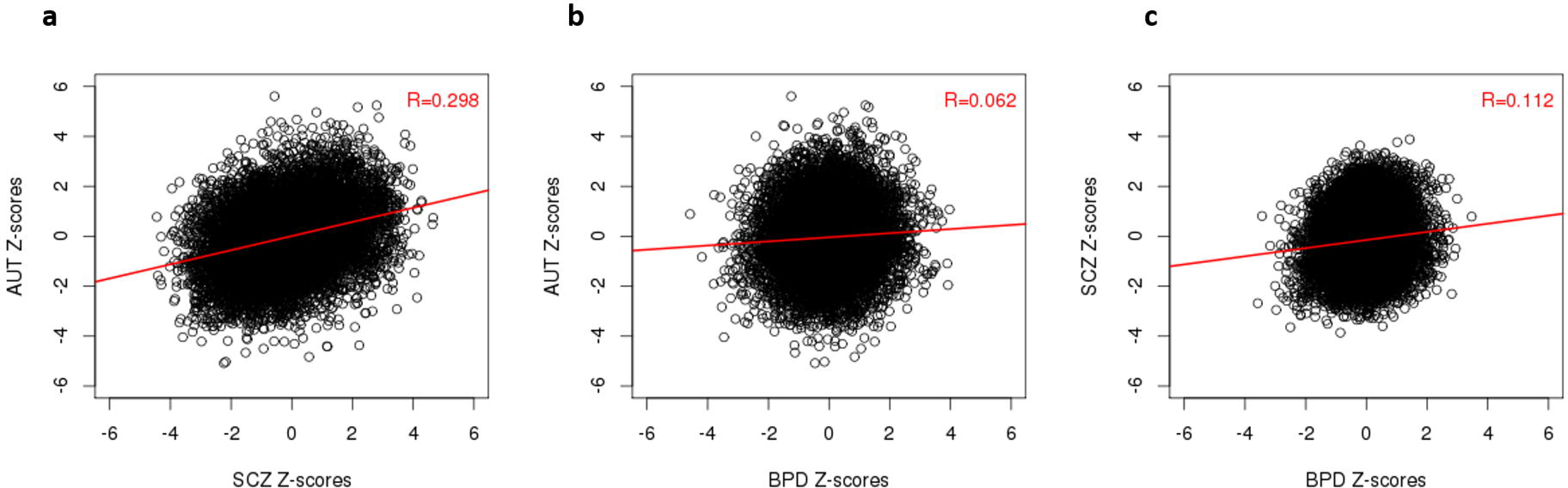
Correlation of Cross-Disorder Differential Gene Expression: Z-scores for each cross-disorder comparison ((a) AUT-SCZ (b) AUT-BPD (c) SCZ-BPD) are plotted. The best fit line is in red. Pearson’s Correlation Coefficient (R) is included on the graph, quantifying the level of correlation between the transcriptomes of each cross-disorder comparison.

To explore the discrepancy between the correlation reported here for SCZ and BPD and that previously reported, we carried out the same analysis without the inclusion of surrogate variables (SVs) in the model. The failure to include unknown covariates in the model led to a marked increase in the correlation between SCZ and BPD (R=0.50), suggesting that the previously reported correlation between these disorders may have been influenced by hidden structure in the data. (Supplemental Figure 4).

### Pathway enrichment analyses of genes differentially expressed across disorders

Combined pathway analysis utilizing lists of genes differentially expressed across disorders (absolute value(Z-score)>2.2 in both disorders) was carried out using both Gene Ontology (GO) enrichment and DAVID pathway analysis. For this analysis, 191 DEGs for AUT-SCZ, 38 for AUT-BPD, and 16 for SCZ-BPD met these criteria. DAVID pathway analysis highlighted the role of neuron projection development (p_Bonferroni_=0.012) in those genes differentially expressed in both AUT and SCZ (Table 3). Similarly, when these genes were characterized by GO, there was a clear abundance of altered gene expression in neuronal and synapse-related GOs (Figure 2). Further, when these DEGS_AUT-SCZ_ were split up into thosegenes were split up into those either concordantly up- or down-regulated in both disorders, 106 genes differentially downregulated in both disorders were driving the GO enrichments, with no contribution from the 69 genes upregulated in both disorders (Supplemental Figure 5). As for AUT-BPD comparisons, there were no enrichments detected for any gene ontologies and the only emergent DAVID pathway was genes related to phosphoproteins (P_Bonferroni_=1-2×10^-4^) (Table 3). Similarly, no GO or DAVD pathways were found to be significant for DEGs_SCZ-BPD_. Substantially similar results were observed when the number of genes from each cross-disorder comparison input into the pathway analysis was fixed rather than imposing a Z-score cutoff (Supplemental Table 5, Supplemental Figures 6,7). Finally, we found that the number of cross-disorder discordant DEGs (upregulated in one disorder but downregulated in the other) differs across the three comparisons, such that there are fewer discordant cross-disorder DEGs (16/191, 8.4%) in the comparison between SCZ and AUT than in the comparison between AUT and BPD (76/191, 39.8%) or between SCZ and BPD (38/191,19.9%), further supporting the transcriptomic similarities between AUT and SCZ.

**Figure 2.**
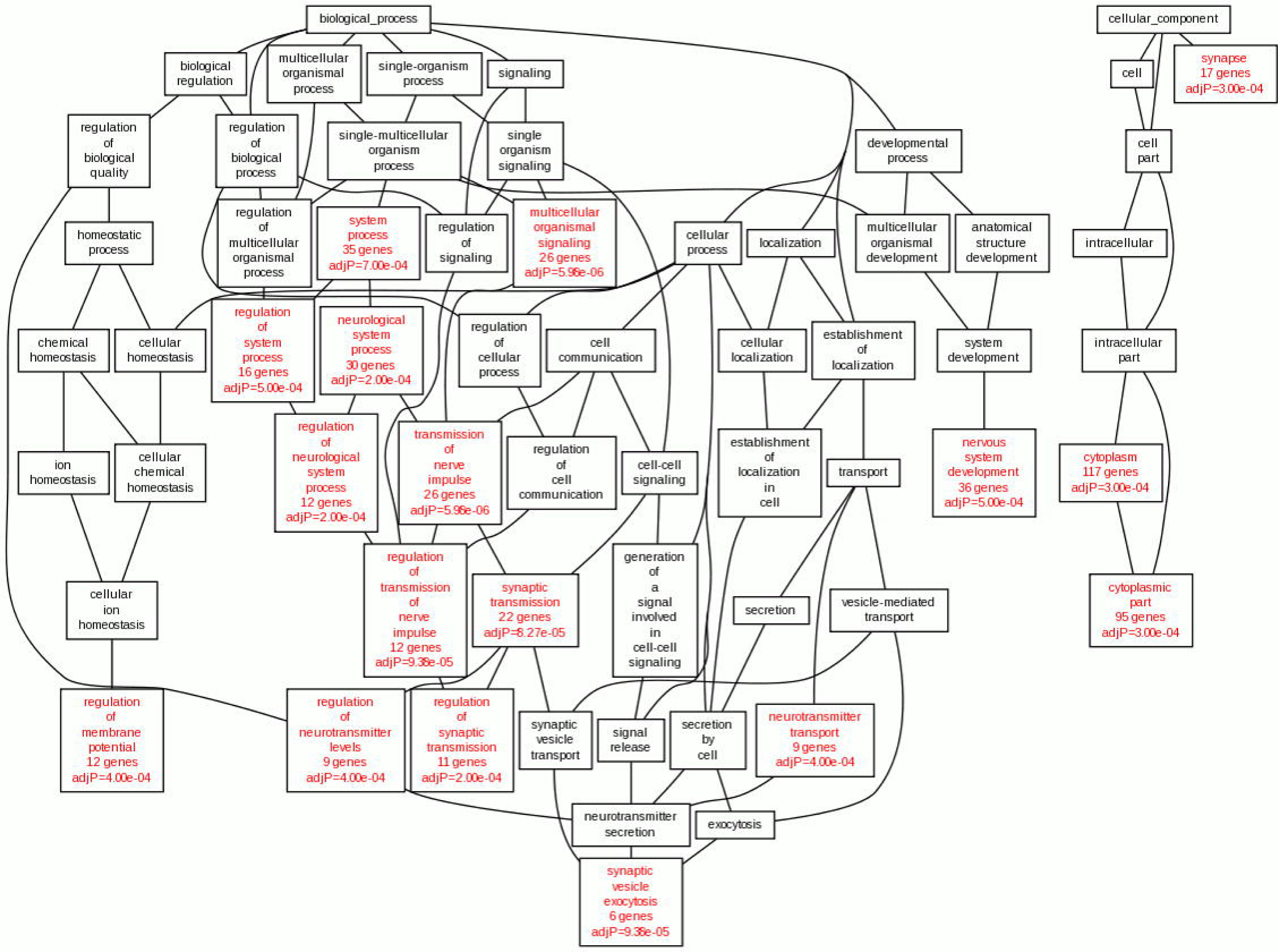
GO Analysis of cross-disorder DEGS_AUT-SCZ_: Genes differentially expressed in both AUT and SCZ (absolute(Z-score) > 2.2) were analyzed for ontological enrichment of biological processes, developmental processes, and cellular component. Onotological categories with at least five genes and an adjusted p-value < 0.001 are highlighted in red. This tree highlights the role of nerve impulse transmission, synaptic transmission, and neurotransmitter transport in those genes differentially expressed in both AUT and SCZ.

**Table 3.** DAVID Pathway Analysis for cross-disorder DEGs. Abbreviations: AUT, autism; BPD, bipolar disorder; SCZ, schizophrenia

|  | TOTAL | UP | DOWN | DISCORDANT | DAVID Pathways |
| --- | --- | --- | --- | --- | --- |
| AUT-SCZ | 191 | 69 | 106 | 0 | neuron projection development (p=0.012) |
| AUT-BPD | 38 | 8 | 19 | 11 | phosphoprotein (p=1.2x10<sup>-4</sup>) |
| SCZ-BPD | 16 | 2 | 13 | 1 | -- |

Abbreviations: AUT, autism; BPD, bipolar disorder; SCZ, schizophrenia

Traditional pathway analysis requires a significance cutoff for the gene input for analysis. To avoid a potential bias by choosing an arbitrary cutoff, we used a Z-score based approach (see Methods) and identified gene enrichment of DCEGs common to all three disorder comparisons using the GO data from MSigDB. Three GO pathways – each of which indicated some enrichment for altered gene expression in transporter genes – were enriched for DEGs in both AUT and SCZ. No pathways were study-wide significant in the other two disorder comparisons (Supplemental Table 6).

### Cross-disorder DEGs enrichment in association signals

To test whether genes differentially expressed across disorders were enriched for genetic associations, we compared cross-disorder DGEA results to gene-level GWAS results. We first directly compared gene-based GWAS p-values (p<0.05) from each individual GWAS (AUT, SCZ, BPD) to p-values from the cross-disorder differential gene expression analysis (AUT-SCZ, AUT-BPD, SCZ-BPD). No comparison was identified that would suggest any enrichment in signal overlap with respect to the null (Supplemental Figure 8). Three additional p-value cutoffs (p<0.1, p<0.01, p<1) demonstrated that neither these null findings nor the inflation seen are a function of the gene-based p-value cutoff imposed on the data (Supplemental Figures 9-11, Supplemental Table 7). Likewise, there were no enrichments for cross-disorder DEGs seen in these analyses relative to the null. Finally, LOF variants have recently been reported in a number of AUT studies (27-32); however, Gupta et. al demonstrated that these gene expression data are neither enriched for the findings from the exome studies nor for Structural Variants (SVs) (12). Accordingly, these lists of variants have not been included in these analyses.

## CONCLUSION AND DISCUSSION

To our knowledge, this is the first study to combine next-generation sequencing gene expression analyses across AUT, SCZ, and BDP to assess the transcriptomic relationship and how gene expression relates to GWAS findings. We report that, at the transcriptome level, AUT and SCZ demonstrate a highly overlapping gene expression profile. The cross-disorder DEGs between AUT and SCZ highlight a shared relationship in synapse and projection formation, suggesting a role for neuronal development underlying the correlation. Further, despite the lack of global significant differential transcriptomic correlation between either BPD and SCZ or AUT and BPD, we highlight two genes, *IQSEC3* and *COPS7A*, for their consistent down regulation across all three disorders and support further investigation into these specific genes’ expression and function to better understand their role in neuropsychiatric disorders. Finally, we report that the genes differentially expressed across disorders were not enriched in genetic association signals for AUT, SCZ or BPD.

### Correlations in Differential Gene Expression Across Disorders Highlights Similarities between AUT and SCZ

After modeling the data for each individual-disorder comparison relative to their controls, the cross-disorder comparison demonstrated that SCZ and AUT share a similarly altered transcriptome (p<0.001), whereas AUT-BPD and SCZ-BPD (p=0.25 and p=0.41, respectively) do not show a significant correlation (Figure 1, Supplemental Figure 3). We note that the lack of significant correlation between BDP-SCZ in our analysis is in conflict with a previous report (10), and is likely due to our inclusion of SVs to account for unknown sources of variation, suggesting that the previously reported analysis of these data is overstated (see Supplemental Discussion). Further, while these data do not directly support transcriptomic overlap between SCZ and BPD, this is likely reflective of the shared control design of the experiment. This experimental design results in a smaller effective sample size and a study underpowered to assess overlap between these two disorders. Given the genetic relationship between these disorders (where SCZ-BPD > AUT-SCZ > AUT-BPD) (1), future work utilizing a larger sample for analysis may likely demonstrate a shared transcriptomic profile between SCZ and BPD; however, these data do not.

Analyzing the pathways in which DEGs in both SCZ and AUT were involved, we found that the genes differentially expressed in AUT and SCZ were enriched for neuron projection development (p=0.012, Table 3). Additionally, there was a clear enrichment for genes involved in synaptic and neuronal processes. The other two non-significant cross-disorder comparisons (AUT-BPD and SCZ-BPD) failed to demonstrate any enrichment for biological process ontology, even when controlling for the number of cross-disorder DEGs, further supporting the conclusion that differential transcriptomic correlation is biologically relevant between SCZ and AUT but is not observed in the other two cross-disorder comparisons (Supplemental Figure 6). When the DEGs across AUT and SCZ were broken down into those concordantly upregulated versus those concordantly downregulated, the enrichment in GO was only present in those genes concordantly downregulated (Supplemental Figure 5), suggesting that these synaptic and neuronal alterations were a result of decreased brain expression in both disorders.

Finally, in assessing which specific genes were differentially expressed across disorders, we identified *IQSEC3* and *COPS7A* as differentially expressed in all three disorders (Table 2). *IQSEC3 (KIAA1110)* is a protein coding gene that has been shown to be specifically expressed in human adult brain with particularly high levels in the human cortex (33). IQSEC3 has been suggested to act as a guanine exchange factor for ARF1 in endocytosis (33), and ARF1 critically regulates actin dynamics in neurons and synaptic strength and plasticity, potentially aligning with pathways previously implicated in AUT, SCZ and BPD. *COPS7A* is expressed broadly across tissues (34), and encodes part of the COP9 signalosome, a multi-subunit protease with a role in regulating the ubiquitin-proteasome pathway (35).

### Differences in Genetic Variation Not Explained by Overlapping Gene Expression Profiles

We report no enrichment for significant cross-disorder DEGs among GWAS signal in any of the comparisons (Supplemental Figure 8) relative to the null. These findings suggest either that 1) alterations at the genetic level do not largely manifest themselves in altered gene expression concordantly across these disorders, or 2) that primary genetic defects do not result in altered gene expression across disorders at the time points measured but could, perhaps, alter gene expression at other time points, such as during development, or 3) the effects of these genetic perturbations are small and that increased sample sizes will be required to detect these slight differences in cross-disorder altered gene expression. Regardless, large differences in gene expression across these disorders appear to be independent of known genetic variation in each of these disorders.

There were a number of limitations associated with our observations. As the analyses combine data across two studies with notable design differences in each (shared controls in the SMRI data, multiple brain regions from the same individual in the AUT data, limited ability to detect lowly expressed genes, and comparison of different cortical brain regions), there was certainly variation unrelated to disease state introduced into the differential gene expression analyses. However, we have controlled for this to the best of our ability by accounting for unknown covariates in all analyses and by determining all levels of significance relative to null permutations. While we have controlled for the differences in experimental design in our analysis, we note that the reported overlap in AUT and SCZ was significant (p<0.001) despite the fact that different cortical brain regions were studied in the two data sets. Due to this limitation, we hypothesize that our observed correlation between AUT and SCZ may underestimate the true transcriptomic correlation and that the similarities may be even more pronounced between AUT and SCZ had the same brain regions been studied. Similarly, sequencing depths in these data sets are lower than many RNA-Seq data sets currently being published. Thus, while lowly-expressed genes are not well-estimated here, their omission from analysis would only lead to false negatives – or genes missing from overlap. This does not detract for the findings, herein, but simply acknowledges that some genes may not be included in the analysis, herein. Conversely, we acknowledge that our power to detect correlation between SCZ and BPD is limited due to the smaller effective sample size, a consequence of the shared control design of the experiment and that, given a larger sample size, transcriptomic correlation between these two disorders may likely become evident and reflective of the known genetic relationships (1).

With future studies employing larger sample sizes and more powerful characterizations, we will gain a better understanding of the transcriptomic relationships that are common and disparate among neuropsychiatric disorders. Besides providing context for how the altered genetic landscape of each disorder affects the brain, we hope that identification of common aspects underlying susceptibility might be novel targets to therapeutically address the underlying pathogenic mechanisms.

## ACKNOWLEDGMENTS

This work was supported by a grant from the Simons Foundation (SFARI 137603 to D.E. Arking) and by the National Institute of Mental Health Autism Centers of Excellence Network Grant 5R01MH081754. We thank Loyal Goff for critical review of the manuscript.

### CONFLICT OF INTEREST

The authors declare no conflict of interest

## SUPPLEMENTAL INFORMATION

Supplemental information is available at *Molecular Psychiatry’s* website

## REFERENCES

1. Cross-Disorder Group of the Psychiatric Genomics Consortium, Lee SH, Ripke S, Neale BM, Faraone SV, Purcell SM, et al. Genetic relationship between five psychiatric disorders estimated from genome-wide SNPs. Nat Genet. 2013 Sep;45(9):984–994.

2. Rutter M. Childhood schizophrenia reconsidered. J Autism Child Schizophr. 1972 Dec; 2(4):315–337.

3. World Health Organization. The ICD-10 Classification of Mental and Behavioural Disorders. American Psychiatric Association; 1993.

4. American Psychiatric Association. Diagnostic and Statistical Manual of Mental Disorders. 4th ed. American Psychiatric Association; 1994.

5. Smoller JW, Finn CT. Family, twin, and adoption studies of bipolar disorder. Am J Med Genet C Semin Med Genet. 2003 Nov 15;123C(1):48–58.

6. Carroll LS, Owen MJ. Genetic overlap between autism, schizophrenia and bipolar disorder. Genome Med. 2009;1(10):102.

7. Ruderfer DM, Fanous AH, Ripke S, McQuillin A, Amdur RL, Schizophrenia Working Group of Psychiatric Genomics Consortium, et al. Polygenic dissection of diagnosis and clinical dimensions of bipolar disorder and schizophrenia. Mol Psychiatry. 2014 Sep;19(9):1017–1024.

8. Cardno AG, Owen MJ. Genetic relationships between schizophrenia, bipolar disorder, and schizoaffective disorder. Schizophr Bull. 2014 May;40(3):504–515.

9. Crespi B, Stead P, Elliot M. Comparative genomics of autism and schizophrenia. Proc Natl Acad Sci. 2010 Jan 26;107(suppl 1):1736–1741.

10. Zhao Z, Xu J, Chen J, Kim S, Reimers M, Bacanu S-A, et al. Transcriptome sequencing and genome-wide association analyses reveal lysosomal function and actin cytoskeleton remodeling in schizophrenia and bipolar disorder. Mol Psychiatry. 2015 May;20(5):563–572.

11. Voineagu I, Wang X, Johnston P, Lowe JK, Tian Y, Horvath S, et al. Transcriptomic analysis of autistic brain reveals convergent molecular pathology. Nature. 2011 Jun 16;474(7351):380–384.

12. Gupta S, Ellis SE, Ashar FN, Moes A, Bader JS, Zhan J, et al. Transcriptome analysis reveals dysregulation of innate immune response genes and neuronal activity-dependent genes in autism. Nat Commun [Internet]. 2014 Dec 10 [cited 2015 Aug 19];5. Available from: <u>http://www.nature.com.ezproxy.welch.jhmi.edu/ncomms/2014/141210/ncomms6748/full/ncomms6748.html</u>

13. Martin M. Cutadapt removes adapter sequences from high-throughput sequencing reads. EMBnetjournal. 2011 Feb 5;17(1):pp. 10–12.

14. Trapnell C, Pachter L, Salzberg SL. TopHat: discovering splice junctions with RNA-Seq. Bioinformatics. 2009 May 1;25(9):1105–1111.

15. Kim D, Pertea G, Trapnell C, Pimentel H, Kelley R, Salzberg SL. TopHat2: accurate alignment of transcriptomes in the presence of insertions, deletions and gene fusions. Genome Biol. 2013;14(4):R36.

16. Hansen KD, Irizarry RA, Wu Z. Removing technical variability in RNA-seq data using conditional quantile normalization. Biostat Oxf Engl. 2012 Apr;13(2):204–216.

17. Ellis SE, Gupta S, Ashar FN, Bader JS, West AB, Arking DE. RNA-Seq optimization with eQTL gold standards. BMC Genomics. 2013 Dec 17;14(1):892.

18. Leek JT, Storey JD. Capturing heterogeneity in gene expression studies by surrogate variable analysis. PLoS Genet. 2007 Sep;3(9):1724–1735.

19. The American Soldier: Vol. I: Adjustment During Army Life, by Samuel A. Stouffer; The American Soldier: Vol. II: Combat and Its Aftermath.

20. WEB-based GEne SeT AnaLysis Toolkit (WebGestalt): update 2013 [Internet], [cited 2015 Aug 25]. Available from: <u>http://nar.oxfordjournals.org/content/41/Wl/W77.full</u>

21. Zhang B, Kirov S, Snoddy J. WebGestalt: an integrated system for exploring gene sets in various biological contexts. Nucleic Acids Res. 2005 Jul 1;33(Web Server issue):W741–W748.

22. Ashburner M, Ball CA, Blake JA, Botstein D, Butler H, Cherry JM, et al. Gene Ontology: tool for the unification of biology. Nat Genet. 2000 May;25(1):25–29.

23. Consortium TGO. Gene Ontology Consortium: going forward. Nucleic Acids Res. 2015 Jan28;43(D1):D1049–D1056.

24. Sherman BT, Huang DW, Tan Q, Guo Y, Bour S, Liu D, et al. DAVID Knowledgebase: a gene-centered database integrating heterogeneous gene annotation resources to facilitate high-throughput gene functional analysis. BMC Bioinformatics. 2007 Nov 2;8(1):426.

25. YH Benjamini YH. Controlling The False Discovery Rate – A Practical And Powerful Approach To Multiple Testing. J R Stat Soc Ser B. 1995;57:289–300.

26. Chanda P, Huang H, Arking DE, Bader JS. Fast association tests for genes with FAST. PloS One. 2013;8(7):e68585.

27. Iossifov I, Ronemus M, Levy D, Wang Z, Hakker I, Rosenbaum J, et al. De Novo Gene Disruptions in Children on the Autistic Spectrum. Neuron. 2012 Apr 26;74(2):285–299.

28. Sanders SJ, Murtha MT, Gupta AR, Murdoch JD, Raubeson MJ, Willsey AJ, et al. De novo mutations revealed by whole-exome sequencing are strongly associated with autism. Nature. 2012 May 10;485(7397):237–241.

29. Neale BM, Kou Y, Liu L, Ma’ayan A, Samocha KE, Sabo A, et al. Patterns and rates of exonic de novo mutations in autism spectrum disorders. Nature [Internet]. 2012 Apr 4 [cited 2012 Apr 13]; Available from: <u>http://www.ncbi.nlm.nih.gov/pubmed/22495311</u>

30. O’Roak BJ, Vives L, Girirajan S, Karakoc E, Krumm N, Coe BP, et al. Sporadic autism exomes reveal a highly interconnected protein network of de novo mutations. Nature [Internet]. 2012 Apr 4 [cited 2012 Apr 13]; Available from: <u>http://www.ncbi.nlm.nih.gov/pubmed/22495309</u>

31. De Rubeis S, He X, Goldberg AP, Poultney CS, Samocha K, Ercument Cicek A, et al. Synaptic, transcriptional and chromatin genes disrupted in autism. Nature. 2014 Nov 13;515(7526):209–215.

32. Talkowski ME, Rosenfeld JA, Blumenthal I, Pillalamarri V, Chiang C, Heilbut A, et al. Sequencing chromosomal abnormalities reveals neurodevelopmental loci that confer risk across diagnostic boundaries. Cell. 2012 Apr 27;149(3):525–537.

33. Hattori Y, Ohta S, Hamada K, Yamada-Okabe H, Kanemura Y, Matsuzaki Y, et al. Identification of a neuron-specific human gene, KIAA1110, that is a guanine nucleotide exchange factor for ARF1. Biochem Biophys Res Commun. 2007 Dec 28;364(4):737–742.

34. Lonsdale J, Thomas J, Salvatore M, Phillips R, Lo E, Shad S, et al. The Genotype-Tissue Expression (GTEx) project. Nat Genet. 2013 Jun;45(6):580–585.

35. Wei N, Serino G, Deng X-W. The COP9 signalosome: more than a protease. Trends Biochem Sci. 2008 Dec;33(12):592–600.

